# The CD112-TIGIT/PVRIG Axis Mediates NK Cell Evasion in Decitabine-Treated Acute Myeloid Leukemia Cells

**DOI:** 10.64898/2026.09.07.749773

**Authors:** Keisuke Nakamura, Ikuko Omori, Naru Sato, Susumu Goyama, Toshio Kitamura, Yutaka Enomoto

**Author notes:** **Corresponding author**: Yutaka Enomoto, Ph.D., Molecular Pharmacology of Malignant Diseases, Graduate School of Pharmaceutical Sciences, The University of Tokyo, 7-3-1 Hongo, Bunkyo-ku, Tokyo 113-0033 Japan.

## Abstract

Both drug resistance and immune evasion are considered to contribute to the persistence of residual AML cells and disease relapse. Natural killer (NK) cells play an important role in antileukemic immune surveillance and represent a key component of antitumor immunity in AML. Here, we investigated whether antileukemic drugs alter the susceptibility of surviving AML cells to natural killer (NK) cell-mediated cytotoxicity. AML cells surviving decitabine treatment became resistant to NK-92-mediated killing and showed marked upregulation of CD112. CD112 overexpression reduced NK-92-mediated cytotoxicity, whereas deletion of CD112 in AML cells or deletion of either TIGIT or PVRIG, two inhibitory receptors for CD112, in NK-92 cells restored cytotoxicity. Mechanistically, decitabine induced demethylation of two enhancer regions within the CD112 locus. Deletion of either enhancer attenuated CD112 upregulation, suggesting that enhancer demethylation contributes to decitabine-induced CD112 expression. Furthermore, antibody-mediated blockade of CD112 or combined blockade of TIGIT and PVRIG enhanced cytotoxicity against decitabine-treated AML cells. CD112 upregulation was also observed following azacitidine or cytarabine treatment, although the magnitude of induction varied across drugs and cell lines. These findings identify CD112 upregulation, which is associated with enhancer demethylation, as a mechanism of drug-induced immune evasion in AML and provide a rationale for combining hypomethylating agents with blockade of the CD112-TIGIT/PVRIG axis to enhance NK cell-mediated antileukemic activity.

## Introduction

Acute myeloid leukemia (AML) is an aggressive hematologic malignancy that remains associated with poor clinical outcomes despite recent therapeutic advances^1^. Although AML occurs across all age groups, its incidence increases markedly with age, with a median age at diagnosis exceeding 65 years^2^. Intensive induction chemotherapy with an anthracycline and cytarabine (Ara-C) (the “7+3” regimen) remains the standard treatment for fit patients, whereas the combination of venetoclax (VEN) and a hypomethylating agent (HMA) is the standard of care for elderly or otherwise unfit patients^1,3,4^. Even after patients achieve complete remission, measurable residual disease (MRD) frequently persists and eventually leads to relapse^5,6^. Because relapsed AML is commonly associated with drug resistance and poor prognosis, novel therapeutic strategies to eradicate MRD and prevent relapse are urgently needed.

Accumulating evidence indicates that AML relapse is driven not only by clonal evolution but also by immune evasion. AML cells evade immune surveillance through multiple mechanisms, including upregulation of immune checkpoint molecules, production of immunosuppressive cytokines, dysregulation of immune-activating ligands, and establishment of an immunosuppressive bone marrow microenvironment^7–13^. These mechanisms contribute to MRD persistence and relapse, highlighting immune evasion as an attractive therapeutic target.

Natural killer (NK) cells are key innate immune effectors that eliminate malignant cells without prior antigen sensitization and play a central role in AML immune surveillance^14,15^. NK cells recognize AML cells through multiple activating receptors and mediate cytotoxicity through perforin/granzyme release and cytokine production^16^. However, AML cells suppress NK cell function by upregulating immune checkpoint molecules, altering activating ligand expression, and remodeling the bone marrow microenvironment^8,10,17,18^. Consequently, restoring NK cell function has attracted increasing attention as a potential therapeutic strategy for AML.

NK cell activity is regulated by the balance between activating and inhibitory receptor signaling^15^. Among these pathways, the DNAM-1 (CD226) axis has been recognized as an important regulator of NK cell function^19^. CD112 (NECTIN2), a shared ligand for the activating receptor DNAM-1 and the inhibitory receptors TIGIT and PVRIG, is expressed on AML cells and on cells from many other tumor types^20–22^. Binding of CD112 to DNAM-1 promotes NK cell activation, whereas its interaction with TIGIT or PVRIG delivers inhibitory signals that attenuate NK cell-mediated cytotoxicity. PVRIG has been reported to bind CD112 with higher affinity than DNAM-1, suggesting that it contributes to the regulation of NK cell activity through the CD112 pathway^23^. Accordingly, the CD112-TIGIT/PVRIG pathway has attracted increasing attention as a potential target for NK cell-based immunotherapy^24–28^. However, the mechanisms regulating CD112 expression in AML cells, particularly in response to anti-leukemic therapy, remain poorly understood.

In this study, we investigated whether anti-leukemic agents modulate the expression of immune regulatory molecules in AML cells. We found that representative anti-leukemic drugs upregulated CD112 expression, thereby suppressing NK cell-mediated cytotoxicity. Blockade of the CD112-TIGIT/PVRIG pathway enhanced NK-92-mediated cytotoxicity. Collectively, our findings identify a previously unrecognized mechanism of drug-induced immune evasion in AML and provide a rationale for combining current anti-leukemic therapies with CD112-TIGIT/PVRIG blockade to improve disease control and prevent relapse.

## Methods

### Cell culture

Human leukemia cell lines NOMO-1 (IFO50474, JCRB Cell Bank), HL-60 (CCL-240, ATCC), and MOLM-13 (ACC-554, DSMZ, Braunschweig, Germany) were cultured in RPMI 1640 medium (Nacalai Tesque) supplemented with 10% fetal bovine serum (FBS; FUJIFILM Wako) and 1% penicillin-streptomycin (Nacalai Tesque). The human NK cell line NK-92 (CRL-2407, ATCC) was cultured in Minimum Essential Medium Alpha (MEMα) supplemented with 12.5% FBS (FUJIFILM Wako), 12.5% horse serum (Gibco), 0.1 mM 2-mercaptoethanol, 0.2 mM myo-inositol (Sigma), 0.02 mM folic acid (Sigma), 200 U/mL human interleukin-2 (IL-2; PeproTech), and 1% penicillin-streptomycin. HEK293T cells (CRL-3216, ATCC) were cultured in Dulbecco’s modified Eagle’s medium (DMEM; Nacalai Tesque) supplemented with 10% FBS and 1% penicillin-streptomycin.

### Flow Cytometric Analysis

Cells were incubated with TruStain FcX Fc Receptor Blocking Solution (BioLegend) for 10 min at 4°C. After Fc receptor blocking, the cells were stained with the indicated fluorophore-conjugated antibodies in FACS buffer for 20 min at 4°C. The cells were then washed with FACS buffer consisting of 2% FBS in PBS and resuspended in FACS buffer containing DAPI. Samples were analyzed using an Attune NxT flow cytometer (Invitrogen), and the data were analyzed using FlowJo software (FlowJo, LLC). The antibodies used in this study are listed in Supplementary Table 1.

### NK-92 Co-culture Assay (Cytotoxicity Assay)

Leukemia cell lines were labeled with 5 μM carboxyfluorescein succinimidyl ester (CFSE; Invitrogen) at room temperature for 5 min. NK-92 cells were co-cultured with CFSE-labeled leukemia cells for 2.5-4 h at effector-to-target (E:T) ratios of 1:1, 4:1, 8:1, and 16:1 in U-bottom 96-well plates. After co-culture, the cells were pelleted by centrifugation, the supernatant was removed, and the cells were resuspended in FACS buffer containing DAPI. The samples were analyzed by flow cytometry. Cytotoxicity was calculated using the following formula: Cytotoxicity (%) = [(experimental lysis (%) - spontaneous lysis (%)) / (100 - spontaneous lysis (%))] × 100.

### Generation of CD112-overexpressing HL-60 cells

A pMYs-IP retroviral vector carrying human CD112 cDNA was used to generate CD112-overexpressing HL-60 cells. The CD112 expression vector was co-transfected with the helper plasmids RD114 and M57 into packaging cells. After 24 h, the medium was replaced, and the cells were incubated for an additional 24 h. The retrovirus-containing supernatants were then collected. For retroviral transduction, polybrene was added to 2 mL of the virus-containing supernatant at a final concentration of 10 μg/mL. HL-60 cells (2.0 × 10^5 cells) were suspended in the virus-containing medium and incubated at 37°C for 4-8 h. Subsequently, 2 mL of fresh culture medium was added, and the cells were cultured for an additional 18 h. The medium was then replaced with fresh culture medium. To establish stable CD112-overexpressing cells, the transduced cells were selected with 1 μg/mL puromycin.

### Gene Knockout Using the CRISPR-Cas9 System

To establish CD112-knockout AML cell lines and TIGIT- or PVRIG-knockout NK-92 cells, we used the CRISPR-Cas9 system. The FUCas9Cherry plasmid (Addgene #70182) was used to generate Cas9-expressing cells. After lentiviral transduction, mCherry-positive cells were sorted using a BD FACSAria III cell sorter (BD Biosciences). The sequence of the TIGIT-targeting sgRNA was obtained from a previous report^22^, whereas the sequences of the other sgRNAs were designed using CHOPCHOP. To generate sgRNA constructs, annealed oligonucleotides were cloned into the lentiGuide-Puro vector (Addgene #52963; RRID: Addgene_52963) at the BsmBI site. Each sgRNA plasmid was co-transfected into HEK293T cells with the lentiviral packaging plasmids psPAX2 (Addgene #12260; RRID: Addgene_12260) and pCMV-VSV-G (Addgene #8454; RRID: Addgene_8454). After 24 h, the medium was replaced, and the cells were incubated for an additional 24-48 h. Lentivirus-containing supernatants were collected and used to transduce Cas9-expressing cells. To obtain knockout cells, the transduced cells were selected with puromycin at 1 μg/mL. The sgRNAs used in this study are listed in Supplementary Table 2.

### Reverse Transcription and Quantitative Real-Time PCR (qPCR)

Total RNA was isolated using TRIzol LS Reagent (Invitrogen). Reverse transcription was performed using ReverTra Ace qPCR Master Mix (TOYOBO) according to the manufacturer’s instructions. Quantitative real-time PCR (qPCR) was performed using SYBR Green qPCR Master Mix (Thermo Fisher Scientific) on a LightCycler 96 system (Roche). GAPDH was used as the reference gene for normalization. Relative mRNA expression levels were calculated using the 2^-ΔΔCt method. All reactions were performed in technical triplicates. The primers used in this study are listed in Supplementary Table 3.

### DNA Methylation Analysis by Bisulfite Sequencing

Total genomic DNA was extracted using the NucleoSpin Tissue Kit (Macherey-Nagel) according to the manufacturer’s instructions. Genomic DNA (240 ng) was subjected to bisulfite conversion using the EZ DNA Methylation Kit (Zymo Research) according to the manufacturer’s instructions. The bisulfite-converted DNA was then subjected to PCR amplification of the target regions. The PCR products were cloned into the pCR-Blunt vector using the Zero Blunt PCR Cloning Kit (Invitrogen) and transformed into *E. coli* DH5α cells. Individual colonies were picked, and plasmid DNA was extracted and sequenced. PCR primers were designed using MethPrimer.

### Re-analysis of Publicly Available RNA-seq Datasets

Publicly available RNA-seq data were obtained from the Gene Expression Omnibus (GEO; accession numbers GSE222822 and GSE240439). Differential gene expression analysis was performed in R using DESeq2. Genes with fewer than 10 total counts across all samples were excluded from the analysis. Volcano plots were generated using ggplot2. Genes were categorized as “Up” when the log2 fold change was > 0.5 and the adjusted P value was < 0.05, as “Down” when the log2 fold change was < −0.5 and the adjusted P value was < 0.05, and as “ns” (not significant) when they did not meet either of these criteria. Adjusted P values were calculated using the Benjamini-Hochberg method, as implemented by default in DESeq2.

### ChIP-Atlas Analysis

Publicly available epigenomic datasets were examined at the CD112 (NECTIN2) locus using the ChIP-Atlas Peak Browser. Genomic profiles of RNA polymerase II, DNase-seq accessibility, H3K4me3, H3K4me1, and H3K27ac were visualized to identify putative promoter and enhancer regions. Candidate regulatory regions were selected based on the overlap of chromatin accessibility and histone modification signals.

### Statistical Analysis

Comparisons between two groups were performed using a two-tailed Student’s t-test. Cytotoxicity assay data were analyzed using two-way repeated-measures ANOVA, followed by a multiple-comparisons test to compare the DMSO control and drug-treated groups at each E:T ratio. Statistical significance is indicated as follows: ns; *P* > 0.05; *, *P* ≤ 0.05; **, *P* ≤ 0.01; ***, *P* ≤ 0.001; ****, *P* ≤ 0.0001, unless stated otherwise. Data are represented as the mean **±** SD, unless otherwise specified. All statistical analyses and data visualizations were performed using GraphPad Prism version 9 (GraphPad Software, RRID:SCR_002798).

## Results

### AML cells surviving decitabine treatment acquire resistance to NK cell-mediated cytotoxicity

To determine whether AML cells surviving anticancer drug treatment become resistant to NK cell-mediated cytotoxicity, NOMO-1, HL-60, and MOLM-13 cells were treated with decitabine, azacitidine, or Ara-C for 48-72 h (Figure 1A). The surviving AML cells were then isolated and co-cultured with NK-92 cells (Supplementary Figure 1A). With the exception of Ara-C-treated HL-60 cells, AML cells that survived drug treatment showed significantly reduced susceptibility to NK-92-mediated cytotoxicity compared with DMSO-treated control cells (Figures 1B and 1C, Supplementary Figure 1B). Among the drugs tested, decitabine produced the greatest reduction in susceptibility to NK-92-mediated cytotoxicity. Given the clinical use of venetoclax in combination with hypomethylating agents for AML, we further examined the effects of combined venetoclax and decitabine treatment^3^. NOMO-1 and HL-60 cells surviving the combination treatment similarly exhibited increased resistance to NK-92-mediated cytotoxicity (Supplementary Figure 1C). Next, we examined whether this resistance persisted after both short-term (48-hour) and long-term (12-day) drug treatment. We found that AML cells that survived long-term decitabine treatment also exhibited significantly greater resistance to NK-92-mediated cytotoxicity than their parental counterparts (Figure 1D, E).

**Figure 1.**
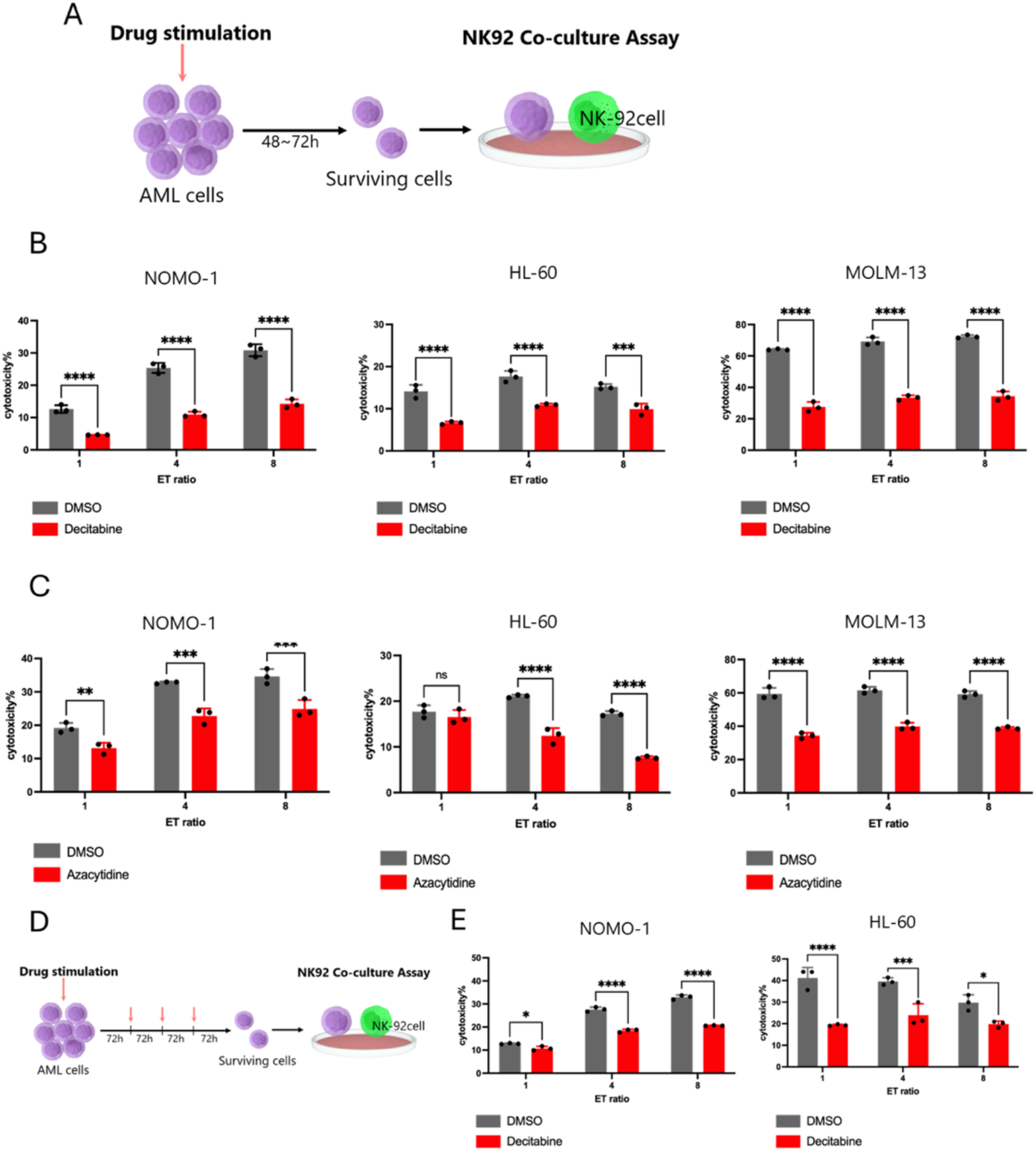
NK-92 cell-mediated cytotoxicity against hypomethylating agent-treated AML cell lines. (A) AML cell lines were treated with AML therapeutic agents. After treatment, the AML cells were collected, washed, and subjected to the NK-92 cytotoxicity assay. (B) NK-92 cell-mediated cytotoxicity against decitabine-treated AML cell lines. NOMO-1, HL-60, and MOLM-13 cells were treated with decitabine under the following conditions: NOMO-1, 1.0 μM for 48 h; HL-60, 3.0 μM for 72 h; and MOLM-13, 1.0 μM for 48 h. After treatment, the AML cells were collected and washed. Target AML cells (5,000 cells/well) were co-cultured with NK-92 cells at the indicated effector-to-target (E:T) ratios for 4 h at 37°C. (C) NOMO-1, HL-60, and MOLM-13 cells were treated with azacitidine under the following conditions: NOMO-1, 1.0 μM for 48 h; HL-60, 1.0 μM for 72 h; and MOLM-13, 2.5 μM for 48 h. After treatment, the AML cells were collected, washed, and subjected to the NK-92 cytotoxicity assay as described in (B). (D and E) NOMO-1 and HL-60 cells were treated with decitabine for 12 days, with the medium replaced every 72 h with fresh decitabine-containing medium. The decitabine concentrations were 0.1 μM for NOMO-1 and 0.3 μM for HL-60. After treatment, the AML cells were collected, washed, and subjected to the NK-92 cytotoxicity assay as described in (B). Data shown are representative of three independent experiments, with each condition assayed in triplicate wells. Data are presented as the mean ± SD. Data were analyzed using two-way ANOVA followed by a multiple-comparisons test to compare the DMSO control and drug-treated groups at each E:T ratio. ns, not significant; *P < 0.05, **P < 0.01, ***P < 0.001, and ****P < 0.0001.

Taken together, these results indicate that AML cells surviving drug treatment acquire resistance to NK cell-mediated cytotoxicity after both short-term and long-term drug treatment.

### Decitabine treatment upregulates CD112 expression on AML cells

We next examined whether decitabine treatment alters the expression of immune regulatory molecules on AML cells (Figure 2A, B, C, Supplementary Figure 2A). Among the molecules examined, CD112 (Nectin-2) surface expression was significantly increased in NOMO-1, MOLM-13, and HL-60 cells following decitabine treatment (Figure 2C, Supplementary Figure 2A). Although the expression of ULBP2/5/6 was also markedly altered following decitabine treatment, these changes were less consistent across the AML cell lines (Supplementary Figure 2A). Furthermore, NOMO-1 cells subjected to long-term decitabine treatment also exhibited increased surface expression of CD112 (Figure 2D).

**Figure 2.**
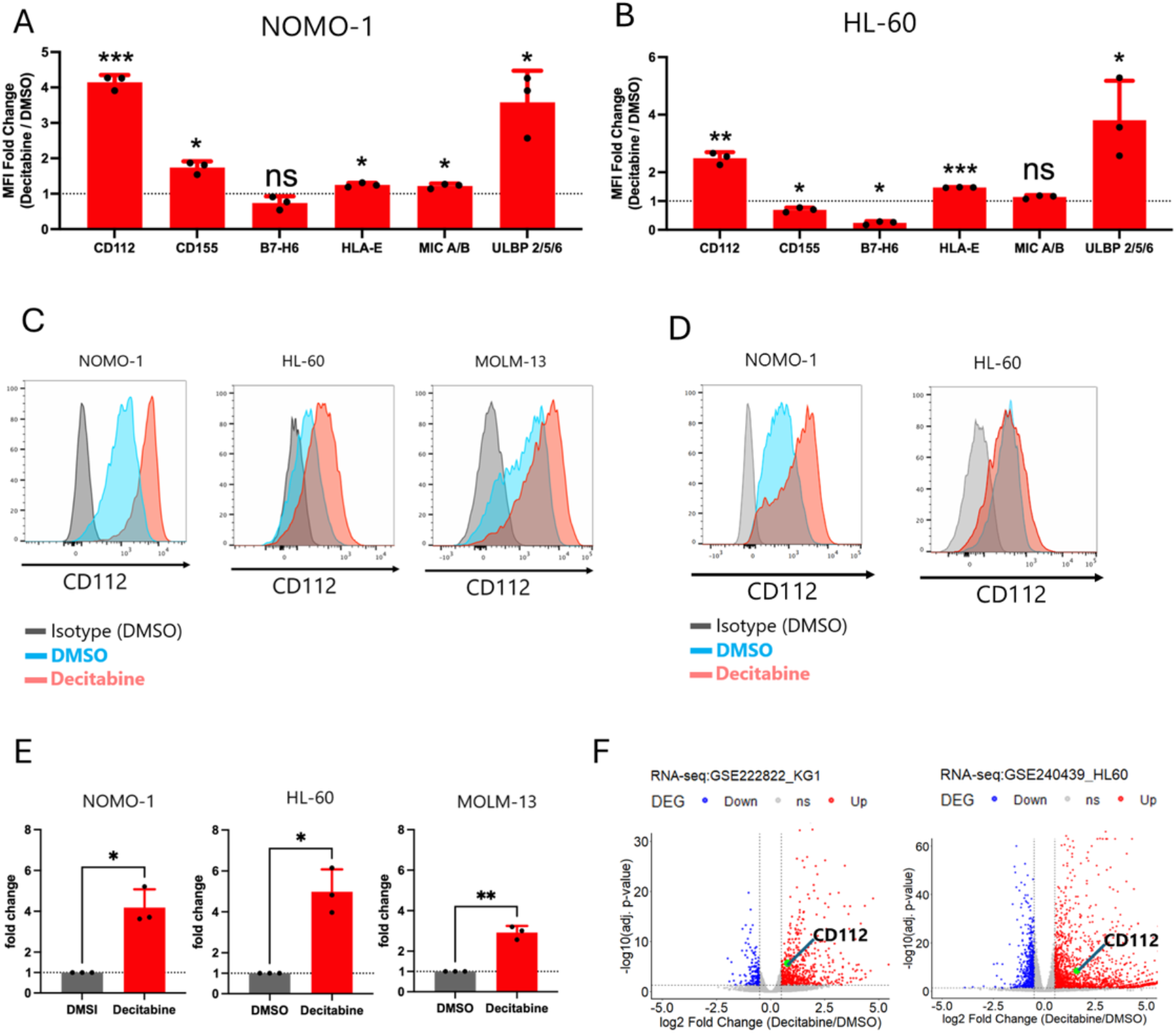
CD112 expression in decitabine-treated AML cells. (A and B) NOMO-1 and HL-60 cells were treated with decitabine under the following conditions: NOMO-1, 1.0 μM for 48 h; and HL-60, 3.0 μM for 72 h. The surface expression of immunoregulatory molecules was analyzed by flow cytometry on NOMO-1 (A) and HL-60 (B) cells. Expression levels are presented as fold changes in mean fluorescence intensity (MFI) relative to those in the corresponding DMSO-treated controls (n = 3). Statistical significance was assessed using a one-sample t-test of log2-transformed fold changes against 0 (corresponding to a fold change of 1). (C) Representative flow cytometry histograms showing CD112 surface expression on NOMO-1, HL-60, and MOLM-13 cells following treatment with decitabine or DMSO. NOMO-1 and HL-60 cells were treated under the conditions described in (A and B), whereas MOLM-13 cells were treated with 1.0 μM decitabine for 48 h. (D) NOMO-1 and HL-60 cells were treated with decitabine for 12 days, with the medium replaced every 72 h with fresh decitabine-containing medium. The decitabine concentrations were 0.1 μM for NOMO-1 and 0.3 μM for HL-60. CD112 surface expression was subsequently analyzed by flow cytometry. (E) NOMO-1, HL-60, and MOLM-13 cells were treated with decitabine or DMSO under the conditions described above, and CD112 mRNA expression was measured by RT-qPCR (n = 3). GAPDH was used as the reference gene. CD112 expression is presented relative to that in the corresponding DMSO-treated controls. (F) Differential gene expression analysis was performed using publicly available RNA-seq datasets of decitabine-treated KG-1 and HL60 cells. The left panel shows the results for decitabine-treated KG-1 cells from GSE222822, and the right panel shows the results for decitabine-treated HL-60 cells from GSE240439. Genes were categorized as Up, Down, or ns according to the criteria described in the Methods. Data shown are representative of three independent experiments. Quantitative data are presented as the mean ± SD. Statistical significance was assessed using a two-tailed Student’s t-test. ns, not significant; *P < 0.05, **P < 0.01, and ***P < 0.001.

Consistent with the increase in surface expression, CD112 mRNA levels were also significantly elevated after decitabine treatment (Figure 2E). Furthermore, reanalysis of publicly available

RNA-seq datasets from KG-1 and HL-60 cells confirmed increased CD112 expression following decitabine treatment (Figure 2F). Similar results were observed with other anticancer agents. Treatment with azacitidine or Ara-C also increased CD112 surface expression on AML cells (Supplementary Figure 2B, C).

We next investigated the expression of CD112-interacting receptors, including PVRIG, TIGIT, and DNAM-1, as well as the activating NK cell receptors NKp46, NKp44, NKp30, and NKG2D, on NK-92 cells. We found that NK-92 cells expressed the inhibitory receptors PVRIG and TIGIT, whereas the activating receptor DNAM-1 was not detectably expressed (Supplementary Figure 2D). In addition, NK-92 cells expressed the activating receptors NKp44 and NKp30 (Supplementary Figure 2D).

These findings suggest that drug-treated AML cells may evade NK cell-mediated cytotoxicity through upregulation of CD112. To directly test this hypothesis, we generated HL-60 cells overexpressing CD112 (Supplementary Figure 2E). CD112 overexpression significantly reduced the susceptibility of HL-60 cells to NK-92 cell-mediated cytotoxicity (Supplementary Figure 2F). Taken together, these results suggest that drug-induced upregulation of CD112 contributes to the resistance of AML cells to NK cell-mediated cytotoxicity.

### Disruption of the CD112-TIGIT/PVRIG signaling axis enhances NK cell-mediated cytotoxicity

To investigate whether disruption of the CD112-TIGIT/PVRIG signaling axis enhances NK cell-mediated cytotoxicity against AML cells, we generated CD112-knockout (KO) NOMO-1 and HL-60 cells (Supplementary Figure 3A, B). We next examined whether CD112-KO cells remained susceptible to NK-92 cell-mediated cytotoxicity following decitabine treatment. We found that decitabine-treated CD112-KO cells remained significantly more susceptible to NK-92 cell-mediated killing than decitabine-treated control cells (Figure 3A, B). In addition, we generated TIGIT-KO and PVRIG-KO NK-92 cells to disrupt CD112-TIGIT/PVRIG signaling (Supplementary Figure 3C). Both TIGIT-KO and PVRIG-KO NK-92 cells exhibited greater cytotoxicity against decitabine-treated NOMO-1 and HL-60 cells than WT NK-92 cells (Figure 3C, D).

**Figure 3.**
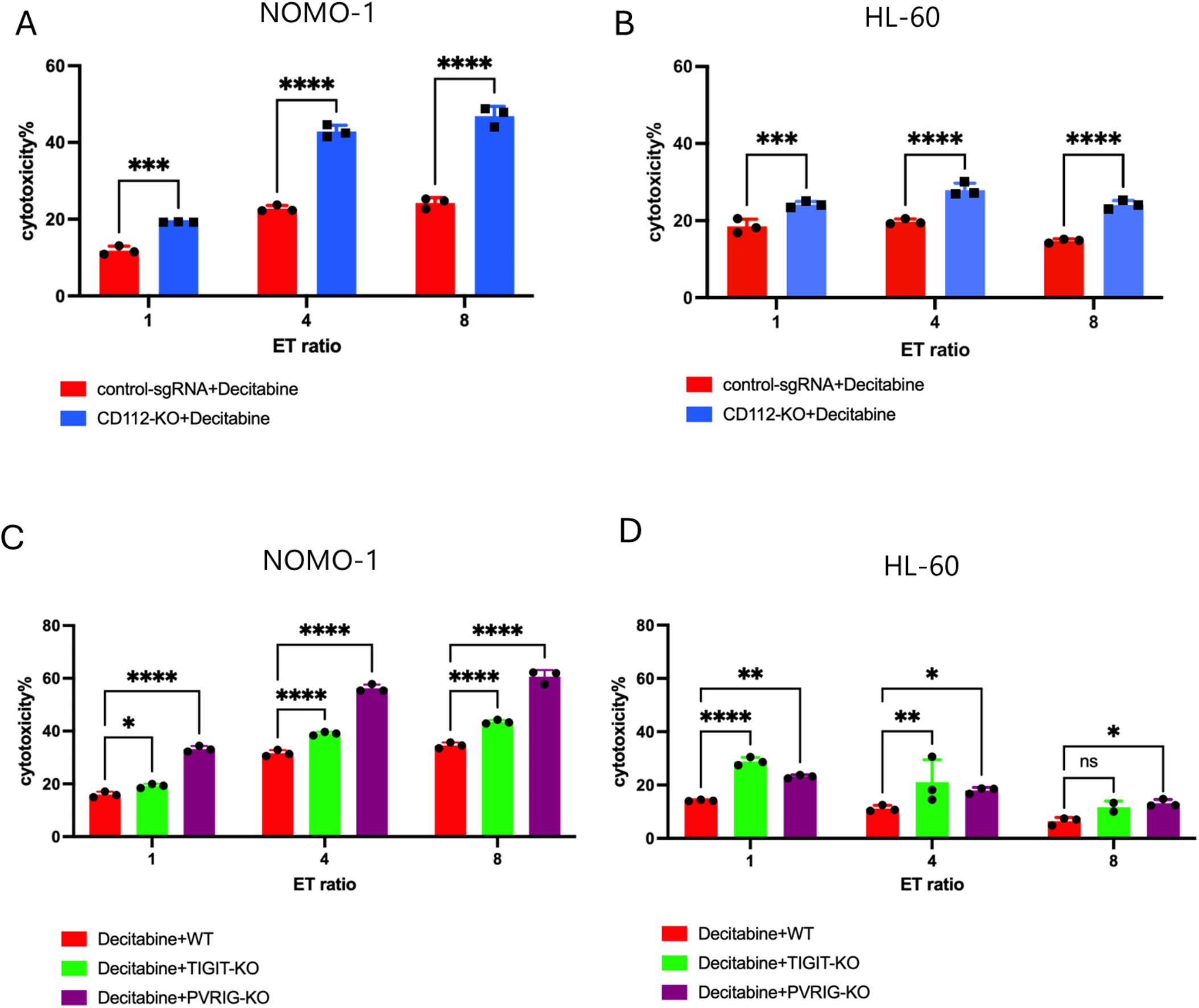
Inhibition of the CD112-inhibitory receptor axis enhances NK-92 cell-mediated cytotoxicity against decitabine-treated AML cells. (A and B) CD112-knockout (CD112-KO) or control sgRNA-expressing NOMO-1 (A) and HL-60 (B) cells were treated with decitabine under the following conditions: NOMO-1, 1.0 μM for 48 h; and HL-60, 3.0 μM for 72 h. After treatment, the AML cells were collected and washed. Target AML cells (10,000 cells/well) were co-cultured with NK-92 cells at the indicated effector-to-target (E:T) ratios for 4 h at 37°C. (C and D) TIGIT-knockout (TIGIT-KO), PVRIG-knockout (PVRIG-KO), or control NK-92 cells were co-cultured with decitabine-treated NOMO-1 (C) and HL-60 (D) cells at the indicated E:T ratios. Cytotoxicity was evaluated after 4 h of co-culture at 37°C. Data were analyzed using two-way ANOVA followed by a multiple-comparisons test to compare the indicated groups at each E:T ratio. Data shown are representative of three independent experiments. ns, not significant; *P < 0.05, **P < 0.01, ***P < 0.001, and ****P < 0.0001.

Taken together, these results indicate that the CD112-TIGIT/PVRIG signaling axis suppresses NK cell-mediated cytotoxicity against decitabine-treated AML cells and suggest that targeting this pathway may enhance NK cell-mediated antitumor immunity.

### Identification of enhancer elements regulating CD112 expression

To investigate the regulatory mechanisms underlying CD112 expression, we analyzed RNA polymerase II binding sites, DNase-seq data, and H3K4me3, H3K4me1, and H3K27ac ChIP-seq datasets using ChIP-Atlas (Figure 4A, Supplementary Figure 4A). Integrative analysis of these datasets identified one putative promoter/enhancer region and two putative enhancer elements within the CD112 locus, designated Enhancer1/Promoter, Enhancer2, and Enhancer3. Given that CD112 expression is markedly upregulated by decitabine treatment, we examined whether the methylation status of these regulatory regions was altered following treatment. We found that Enhancer2 and Enhancer3 underwent significant demethylation after decitabine treatment (Figure 4B, C), whereas the methylation status of the Enhancer1/Promoter region remained unchanged (Supplementary Figure 4B). Similarly, azacitidine treatment induced demethylation of Enhancer2, whereas Ara-C treatment did not alter its methylation status (Supplementary Figure 4C, D).

**Figure 4.**
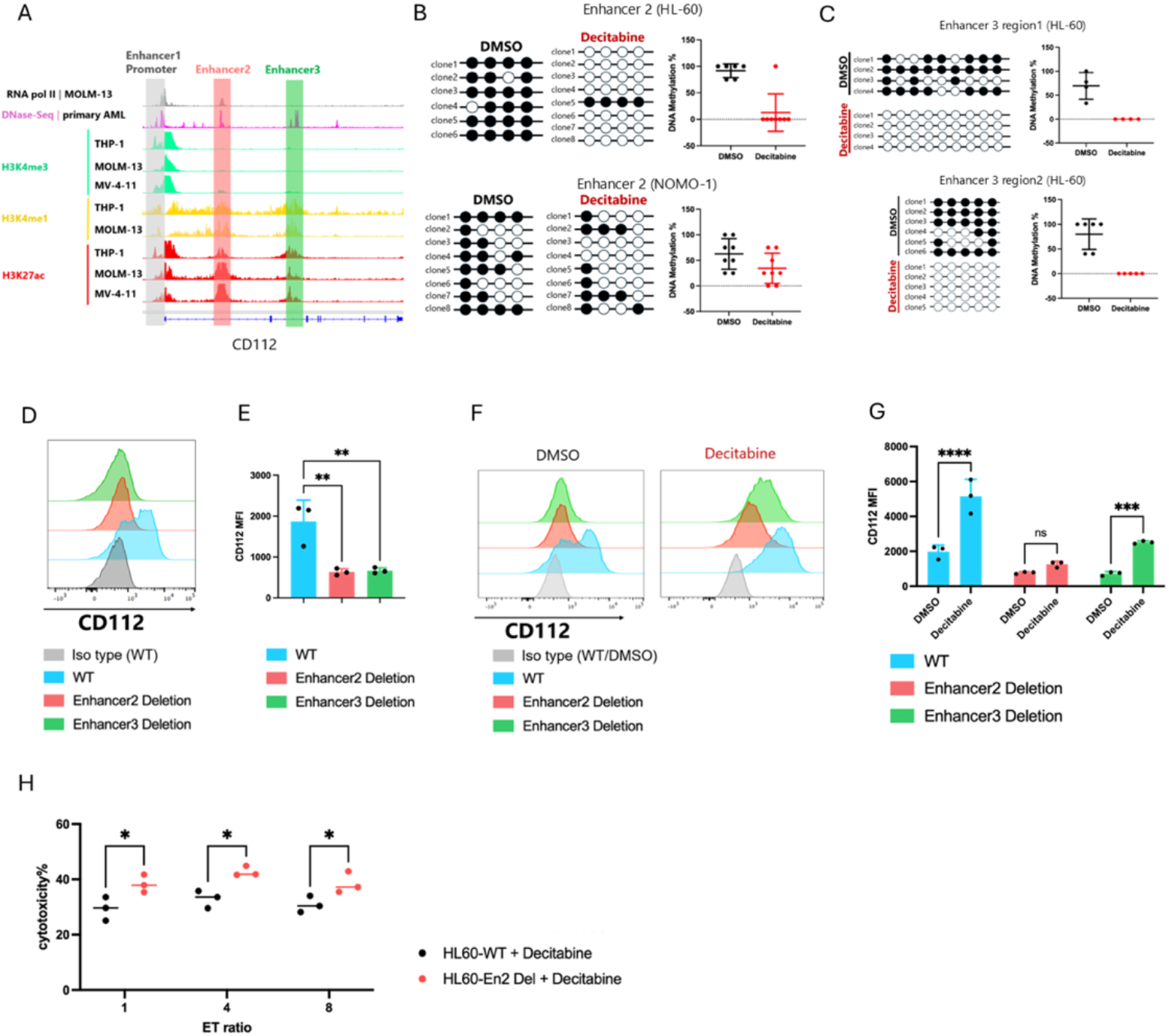
Identification of CD112 regulatory elements involved in decitabine-induced CD112 upregulation. (A) Identification of candidate regulatory elements within the CD112 locus using publicly available epigenomic datasets. ChIP-seq and DNase-seq datasets obtained from ChIP-Atlas were visualized using the Integrative Genomics Viewer (IGV). Genomic tracks for RNA polymerase II (RNA Pol II), DNase I hypersensitivity, H3K4me3, H3K4me1, and H3K27ac across the CD112 locus are shown for primary AML samples and AML cell lines, including THP-1, MOLM-13, and MV4-11. (B) HL-60 cells were treated with 3.0 μM decitabine for 72 h, and NOMO-1 cells were treated with 1.0 μM decitabine for 48 h. Genomic DNA from each sample was subjected to bisulfite sequencing analysis of the CD112 enhancer 2 (En2) region. Representative lollipop plots show the methylation status of individual CpG sites in each clone. Open and filled circles indicate unmethylated and methylated CpG sites, respectively. The percentage of methylated CpG sites was calculated from the bisulfite sequencing results and is shown in the right panels. (C) HL-60 cells were treated with 3.0 μM decitabine or DMSO for 72 h, and genomic DNA was subjected to bisulfite sequencing analysis of two regions within CD112 enhancer 3 (En3 regions 1 and 2). (D and E) CD112 surface expression on wild-type (WT) HL-60 cells and HL-60 cells carrying deletions of enhancer 2 (En2) or enhancer 3 (En3) was analyzed by flow cytometry. Representative histograms are shown in (D), and quantification of CD112 mean fluorescence intensity (MFI) is shown in (E) (n = 3). (F and G) WT, En2-deleted, and En3-deleted HL-60 cells were treated with 3.0 μM decitabine or DMSO for 72 h, and CD112 surface expression was subsequently analyzed by flow cytometry. Representative histograms are shown in F, and quantification of CD112 MFI is shown in G (n = 3). (H) WT and En2-deleted HL-60 target cells (10,000 cells/well) were co-cultured with NK-92 cells at the indicated effector-to-target (E:T) ratios for 4 h at 37°C, followed by evaluation of NK-92 cell-mediated cytotoxicity. Quantitative data are presented as he mean ± SD. Statistical analyses were performed using one-way or two-way ANOVA followed by a multiple-comparisons test, as appropriate. Data shown are representative of three independent experiments. ns, not significant; *P < 0.05, **P < 0.01, ***P < 0.001, and ****P < 0.0001.

To determine whether Enhancer2 and Enhancer3 regulate CD112 expression, we generated cells in which either Enhancer2 or Enhancer3 had been individually deleted (Supplementary Figure 5A-C). Deletion of either enhancer reduced CD112 expression under steady-state conditions and attenuated decitabine-induced CD112 upregulation (Figure 4D-G). Notably, deletion of Enhancer2 had a greater effect on CD112 expression than deletion of Enhancer3. Given the stronger effect of Enhancer2 deletion, we focused on this enhancer in subsequent functional analyses. HL-60 cells carrying an Enhancer2 deletion were significantly more susceptible to NK-92 cell-mediated cytotoxicity than parental cells (Figure 4H).

Taken together, these results suggest that Enhancer2 and Enhancer3 regulate CD112 expression and that decitabine-induced demethylation of these enhancer regions contributes to the induction of CD112 expression.

### Blocking antibodies against the CD112-TIGIT/PVRIG signaling axis enhance NK cell-mediated cytotoxicity

Our findings indicate that drug-induced upregulation of CD112 suppresses NK cell-mediated cytotoxicity against AML cells through the CD112-TIGIT/PVRIG signaling axis. Therefore, to explore the translational potential of targeting this pathway, we evaluated whether blocking antibodies against CD112, TIGIT, and PVRIG could enhance NK cell-mediated cytotoxicity (Figure 5A). Treatment with an anti-CD112 antibody significantly enhanced NK-92 cell-mediated cytotoxicity against decitabine-treated NOMO-1 cells (Figure 5B). Similarly, combined blockade of TIGIT and PVRIG significantly enhanced NK-92 cell-mediated cytotoxicity against decitabine-treated NOMO-1 cells (Figure 5C).

**Figure 5.**
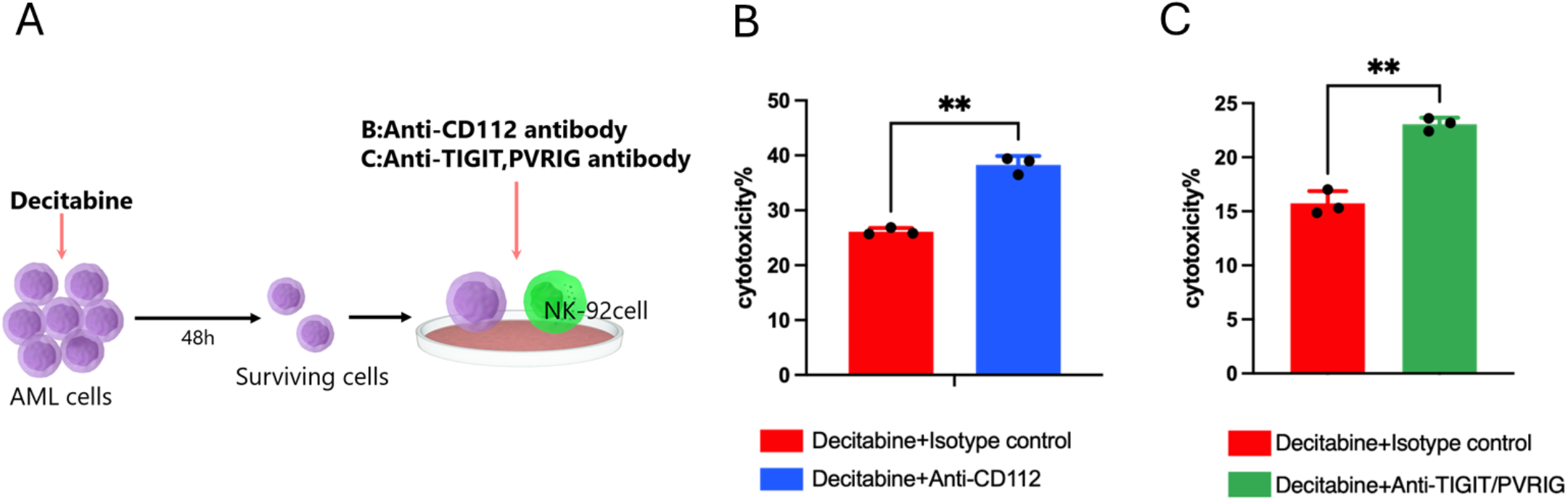
Effects of blocking antibodies targeting the CD112-TIGIT/PVRIG axis on NK-92 cell-mediated cytotoxicity against decitabine-treated AML cells. (A) NOMO-1 cells were treated with 1.0 μM decitabine for 48 h. After treatment, the cells were collected, washed, and used as target cells in the NK-92 cytotoxicity assay. An anti-CD112 antibody (final concentration, 10 μg/mL) was added to the cytotoxicity assay in (B), whereas anti-TIGIT and anti-PVRIG antibodies (final concentration, 40 μg/mL each) were added in (C). (B) Following decitabine treatment, NOMO-1 cells were collected and washed. Target cells (10,000 cells/well) were co-cultured with NK-92 cells at an effector-to-target (E:T) ratio of 4:1 for 4 h at 37°C in the presence or absence of an anti-CD112 antibody (10 μg/mL). (C) Following decitabine treatment, NOMO-1 cells were collected and washed. Target cells (10,000 cells/well) were co-cultured with NK-92 cells at an effector-to-target (E:T) ratio of 2:1 for 4 h at 37°C in the presence or absence of anti-TIGIT and anti-PVRIG antibodies (40 μg/mL each). Data shown are representative of three independent experiments. **P < 0.01.

Taken together, these results suggest that blockade of the CD112-TIGIT/PVRIG signaling axis using antibodies against CD112, TIGIT, and/or PVRIG may represent a promising therapeutic strategy to enhance NK cell-mediated antitumor immunity against AML cells that survive anticancer drug treatment.

## Discussion

In the present study, we demonstrated that AML cells surviving decitabine treatment acquire resistance to NK cell-mediated cytotoxicity and identified CD112 upregulation as one of the major mechanisms underlying this phenotype. We further showed that decitabine-induced CD112 expression is regulated through enhancer demethylation and that genetic or antibody-mediated blockade of the CD112-TIGIT/PVRIG signaling axis restores NK cell-mediated cytotoxicity. Although increased CD112 expression was also observed under certain conditions following azacitidine or Ara-C treatment, the extent of induction varied among drugs and cell lines, suggesting that this phenomenon is not unique to decitabine but may depend on both the therapeutic agent and cellular context. Collectively, these findings suggest that anti-leukemic therapies not only exert direct cytotoxic effects on AML cells but may also induce an adaptive immune evasion program in the cells that survive treatment.

Hypomethylating agents (HMAs) have recently attracted considerable attention because of their ability to enhance anti-tumor immunity through epigenetic reprogramming^29–31^. HMAs have been reported to promote anti-tumor immunity through multiple mechanisms, including viral mimicry, activation of type I interferon signaling, and enhanced antigen presentation^32–34^. Conversely, HMAs have also been shown to induce the expression of immune regulatory molecules, including immune checkpoint molecules, suggesting that they exert both immunostimulatory and immunosuppressive effects^35–37^. In the present study, we identified CD112 as another immune regulatory molecule induced by decitabine.

CD112 plays a complex role in NK cell biology because it serves as a shared ligand for the activating receptor DNAM-1 as well as the inhibitory receptors TIGIT and PVRIG^25^. However, these receptors do not bind CD112 with equal affinity. Previous studies have demonstrated that PVRIG binds CD112 with higher affinity than DNAM-1, whereas TIGIT, although capable of binding CD112, preferentially recognizes CD155 with substantially higher affinity^23,25^. These observations raise the possibility that HMA-induced CD112 expression may function as a novel immune checkpoint mechanism through the PVRIG-CD112 axis, depending on the receptor-expression profile of the NK cells.

In our experimental system, NK-92 cells expressed TIGIT and PVRIG but exhibited little detectable DNAM-1 expression. Therefore, decitabine-induced CD112 expression is likely to engage predominantly the inhibitory receptors TIGIT and PVRIG rather than the activating receptor DNAM-1. Consistent with this interpretation, CD112 overexpression reduced NK cell-mediated cytotoxicity, whereas genetic disruption of CD112, TIGIT, or PVRIG restored NK cell cytotoxicity. These findings support the notion that decitabine-induced CD112 expression shifts the balance toward inhibitory signaling, thereby promoting immune evasion by residual AML cells. Nevertheless, because primary NK cells expressing physiological levels of DNAM-1 were not examined in the present study, the relative contributions of DNAM-1, TIGIT, and PVRIG under physiological conditions remain to be determined.

The concept of therapy-induced immune evasion proposed in this study may have important implications for understanding AML relapse. Despite achieving complete remission, many AML patients retain measurable residual disease (MRD), which ultimately leads to disease recurrence^1^. Our findings suggest that AML cells surviving decitabine treatment may evade NK cell-mediated immune surveillance through CD112 upregulation, thereby contributing to MRD persistence and relapse. Accordingly, targeting the CD112-TIGIT/PVRIG signaling axis in combination with HMA therapy may facilitate more efficient elimination of residual leukemic cells. Indeed, blockade of CD112, TIGIT, or PVRIG restored NK cell-mediated cytotoxicity in our experimental models, supporting the potential of this pathway as a therapeutic target for preventing AML relapse.

This study has several limitations. First, our functional analyses were performed primarily using AML cell lines and the NK-92 cell line, and therefore require validation using primary AML samples and primary human NK cells. Second, because NK-92 cells expressed little DNAM-1, we were unable to fully evaluate the balance between DNAM-1-mediated activating signals and TIGIT/PVRIG-mediated inhibitory signals following CD112 induction. Third, although we demonstrated that enhancer demethylation represents a major mechanism responsible for decitabine-induced CD112 expression, additional upstream transcription factors and chromatin regulatory mechanisms remain to be elucidated. Furthermore, CD112 induction by azacitidine and Ara-C was not consistently observed across all AML cell lines, suggesting that both drug-specific and cell-intrinsic factors influence this response. Finally, the therapeutic efficacy of combining HMA treatment with blockade of the CD112-TIGIT/PVRIG axis should be validated in immunocompetent AML models and patient-derived samples.

In conclusion, we demonstrate that decitabine induces CD112 expression through enhancer demethylation, thereby promoting evasion of NK cell-mediated cytotoxicity via the CD112-TIGIT/PVRIG signaling axis. These findings provide new insights into the interplay between epigenetic therapy and innate immunity and identify therapy-induced immune evasion as a previously unrecognized mechanism that may contribute to AML relapse. Furthermore, our study suggests that targeting the CD112-TIGIT/PVRIG pathway in combination with HMA therapy represents a promising strategy to enhance the elimination of residual leukemic cells and reduce disease recurrence.

## Acknowledgments

This work was supported by JSPS KAKENHI Grant Number 22K08496 and research grants from Japan Leukemia Research Fund (JLRF), Yakugaku Shinkokai Foundation, and International Joint Usage/Research Center, the Institute of Medical Science, the University of Tokyo. K.N. gratefully acknowledges support from ST SPRING (SPRING GX), Grant Number JPMJSP2108.

## Authorship Contributions

K.N. designed and performed most of the experiments, analyzed and interpreted the data, and wrote the manuscript; I.O. and N.S. advised on data interpretation; S.G. and T.K. conceived the project and interpreted the data; Y.E. supervised and conceived the project, designed the experiments, interpreted the data, and wrote the manuscript. All authors critically reviewed and edited the manuscript.

## References

1. Kantarjian, H. et al. Acute myeloid leukemia: current progress and future directions. Blood Cancer J. 11, 41 (2021).

2. Khwaja, A., et al. Acute myeloid leukaemia. Nat. Rev. Dis. Primer 2, 16010 (2016).

3. DiNardo, C. D. et al. Azacitidine and Venetoclax in Previously Untreated Acute Myeloid Leukemia. N. Engl. J. Med. 383, 617–629 (2020).

4. Kwag, D. et al. Venetoclax with decitabine versus decitabine monotherapy in elderly acute myeloid leukemia: a propensity score-matched analysis. Blood Cancer J. 12, 169 (2022).

5. Stelmach, P. & Trumpp, A. Leukemic stem cells and therapy resistance in acute myeloid leukemia. Haematologica 108, 353–366 (2023).

6. Short, N. J. et al. Advances in the Treatment of Acute Myeloid Leukemia: New Drugs and New Challenges. Cancer Discov. 10, 506–525 (2020).

7. Christopher, M. J. et al. Immune Escape of Relapsed AML Cells after Allogeneic Transplantation. N. Engl. J. Med. 379, 2330–2341 (2018).

8. Paczulla, A. M., et al. Absence of NKG2D ligands defines leukaemia stem cells and mediates their immune evasion. Nature 572, 254–259 (2019).

9. Kumar, B. et al. BATF is a major driver of NK cell epigenetic reprogramming and dysfunction in AML. Sci. Transl. Med. 16, eadp0004 (2024).

10. Wang, D. et al. GARP-mediated active TGF-β1 induces bone marrow NK cell dysfunction in AML patients with early relapse post-allo-HSCT. Blood 140, 2788–2804 (2022).

11. Mumme, H. et al. Single-cell analysis reveals altered tumor microenvironments of relapse- and remission-associated pediatric acute myeloid leukemia. Nat. Commun. 14, 6209 (2023).

12. Corradi, G. et al. Release of IFN γ by Acute Myeloid Leukemia Cells Remodels Bone Marrow Immune Microenvironment by Inducing Regulatory T Cells. Clin. Cancer Res. 28, 3141–3155 (2022).

13. Noviello, M. et al. Bone marrow central memory and memory stem T-cell exhaustion in AML patients relapsing after HSCT. Nat. Commun. 10, 1065 (2019).

14. Ruggeri, L. et al. Effectiveness of Donor Natural Killer Cell Alloreactivity in Mismatched Hematopoietic Transplants. Science 295, 2097–2100 (2002).

15. Wolf, N. K., Kissiov, D. U. & Raulet, D. H. Roles of natural killer cells in immunity to cancer, and applications to immunotherapy. Nat. Rev. Immunol. 23, 90–105 (2023).

16. Baragaño Raneros, A., López-Larrea, C. & Suárez-Álvarez, B. Acute myeloid leukemia and NK cells: two warriors confront each other. OncoImmunology 8, e1539617 (2019).

17. Xie, J. et al. Overexpressing natural killer group 2 member A drives natural killer cell exhaustion in relapsed acute myeloid leukemia. Signal Transduct. Target. Ther. 10, 143 (2025).

18. Chang, Y.-H. et al. SETDB1 suppresses NK cell-mediated immunosurveillance in acute myeloid leukemia with granulo-monocytic differentiation. Cell Rep. 43, 114536 (2024).

19. Alteber, Z., et al. Therapeutic Targeting of Checkpoint Receptors within the DNAM1 Axis. Cancer Discov. 11, 1040–1051 (2021).

20. Stamm, H. et al. Immune checkpoints PVR and PVRL2 are prognostic markers in AML and their blockade represents a new therapeutic option. Oncogene 37, 5269–5280 (2018).

21. Kaito, Y. et al. CD155 and CD112 as possible therapeutic targets of *FLT3* inhibitors for acute myeloid leukemia. Oncol. Lett. 23, 51 (2022).

22. Kaito, Y. et al. Immune checkpoint molecule DNAM-1/CD112 axis is a novel target for NK-cell therapy in acute myeloid leukemia. Haematologica 109(4):1107–1120 (2024)

23. Zhu, Y. et al. Identification of CD112R as a novel checkpoint for human T cells. J. Exp. Med. 213, 167–176 (2016).

24. Xu, F. et al. Blockade of CD112R and TIGIT signaling sensitizes human natural killer cell functions. Cancer Immunol. Immunother. 66, 1367–1375 (2017).

25. Zeng, T. et al. The CD112R/CD112 axis: a breakthrough in cancer immunotherapy. J. Exp. Clin. Cancer Res. 40, 285 (2021).

26. Whelan, S. et al. PVRIG and PVRL2 Are Induced in Cancer and Inhibit CD8+ T-cell Function. Cancer Immunol. Res. 7, 257–268 (2019).

27. Li, J. et al. PVRIG is a novel natural killer cell immune checkpoint receptor in acute myeloid leukemia. Haematologica 106, 3115–3124 (2021).

28. Lin, Y. et al. Co-blocking TIGIT and PVRIG Using a Novel Bispecific Antibody Enhances Antitumor Immunity. Mol. Cancer Ther. 24, 664–677 (2025).

29. Daver, N. et al. Hypomethylating agents in combination with immune checkpoint inhibitors in acute myeloid leukemia and myelodysplastic syndromes. Leukemia 32, 1094–1105 (2018).

30. Kang, S. et al. Decitabine enhances targeting of AML cells by NY-ESO-1-specific TCR-T cells and promotes the maintenance of effector function and the memory phenotype. Oncogene 41, 4696–4708 (2022).

31. Li, Y.-R. et al. Allogeneic CD33-directed CAR-NKT cells for the treatment of bone marrow-resident myeloid malignancies. Nat. Commun. 16, 1248 (2025).

32. Huang, C. et al. Priming with DNMT Inhibitors Potentiates PD-1 Immunotherapy by Triggering Viral Mimicry in Relapsed/Refractory NK/T-cell Lymphoma. Cancer Discov. 15, 2450–2467 (2025).

33. Lee, J. B. et al. Venetoclax enhances T cell-mediated antileukemic activity by increasing ROS production. Blood 138, 234–245 (2021).

34. Chiappinelli, K. B. et al. Inhibiting DNA Methylation Causes an Interferon Response in Cancer via dsRNA Including Endogenous Retroviruses. Cell 162, 974–986 (2015).

35. Ørskov, A. D., et al. Hypomethylation and up-regulation of *PD-1* in T cells by azacytidine in MDS/AML patients: A rationale for combined targeting of PD-1 and DNA methylation. Oncotarget 6, 9612–9626 (2015).

36. Yang, H. et al. Expression of PD-L1, PD-L2, PD-1 and CTLA4 in myelodysplastic syndromes is enhanced by treatment with hypomethylating agents. Leukemia 28, 1280–1288 (2014).

37. Guo, H.-Z. et al. A CD36-dependent non-canonical lipid metabolism program promotes immune escape and resistance to hypomethylating agent therapy in AML. Cell Rep. Med. 5, 101592 (2024).

