## Supplementary Figures and Tables for "The CD112-TIGIT/PVRIG Axis Mediates NK Cell Evasion in Decitabine-Treated Acute Myeloid Leukemia Cells"

### Supplementary Figure 1

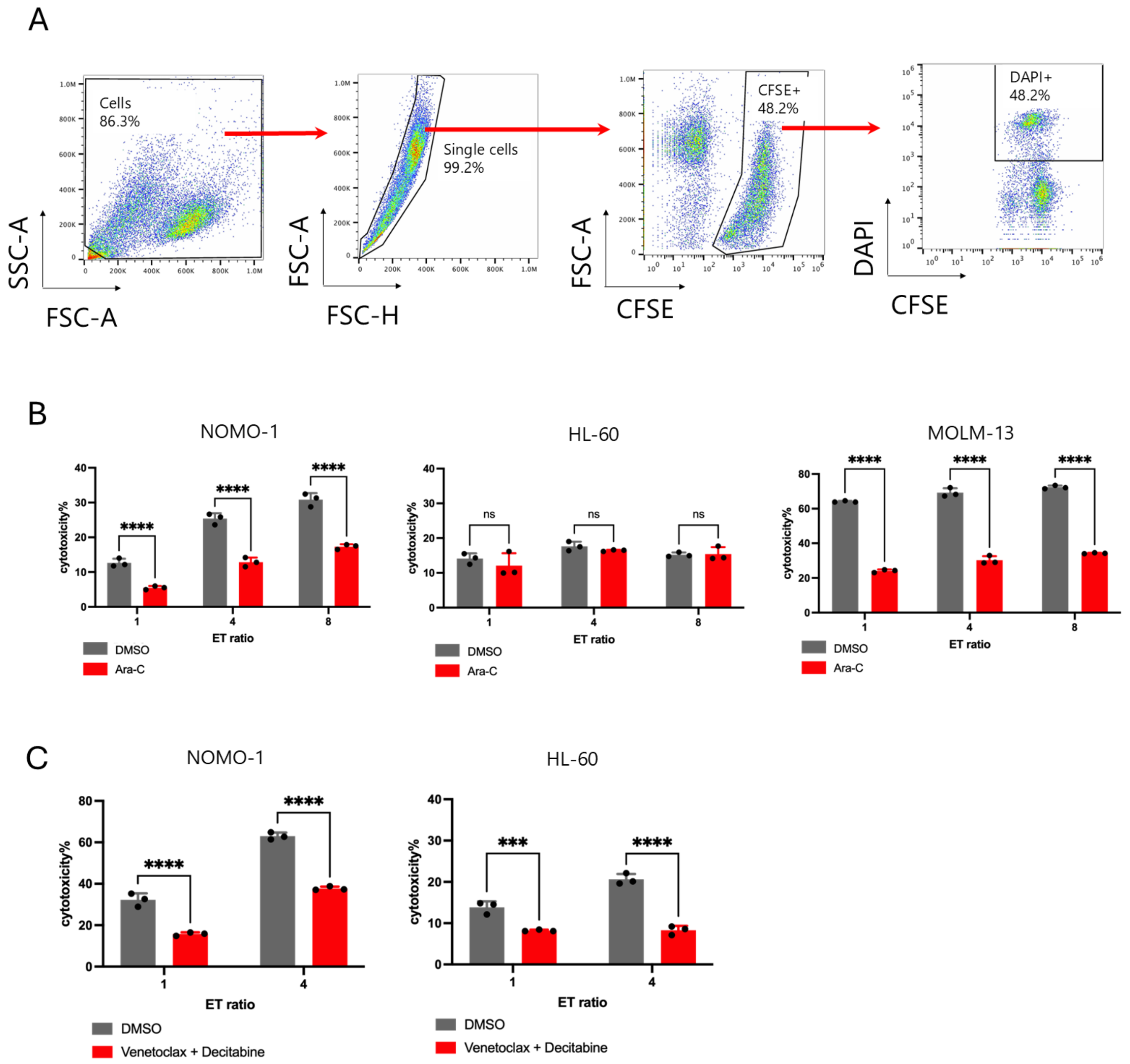

**Supplementary Figure 1. NK-92 cytotoxicity assay and cytotoxicity against drug-treated AML cells**

(A) The gating strategy used to identify DAPI-positive target cells is shown. Flow cytometry data from the NK-92 cytotoxicity assay were analyzed using FlowJo software. Cytotoxicity was calculated based on the proportion of DAPI-positive target cells. (B) NOMO-1, HL-60, and MOLM-13 cells were treated with DMSO or Ara-C under the following conditions: NOMO-1, 1.0 μM for 48 h; HL-60, 3.0 μM for 72 h; and MOLM-13, 0.01 μM for 48 h. After treatment, the AML cells were collected, washed, and used as target cells in the NK-92 cytotoxicity assay. (C) NOMO-1 and HL-60 cells were treated with DMSO or a combination of venetoclax and decitabine under the following conditions: NOMO-1, 1.0 μM venetoclax plus 1.0 μM decitabine for 48 h; and HL-60, 1.0 μM venetoclax plus 3.0 μM decitabine for 72 h. After treatment, the AML cells were collected, washed, and used as target cells in the NK-92 cytotoxicity assay. Data are representative of three independent experiments and are presented as the mean ± SD of triplicate measurements. Data were analyzed using two-way ANOVA followed by a multiple-comparisons test. ns, not significant; ***$P < 0.001$ and ****$P < 0.0001$.

**Supplementary Figure 2**

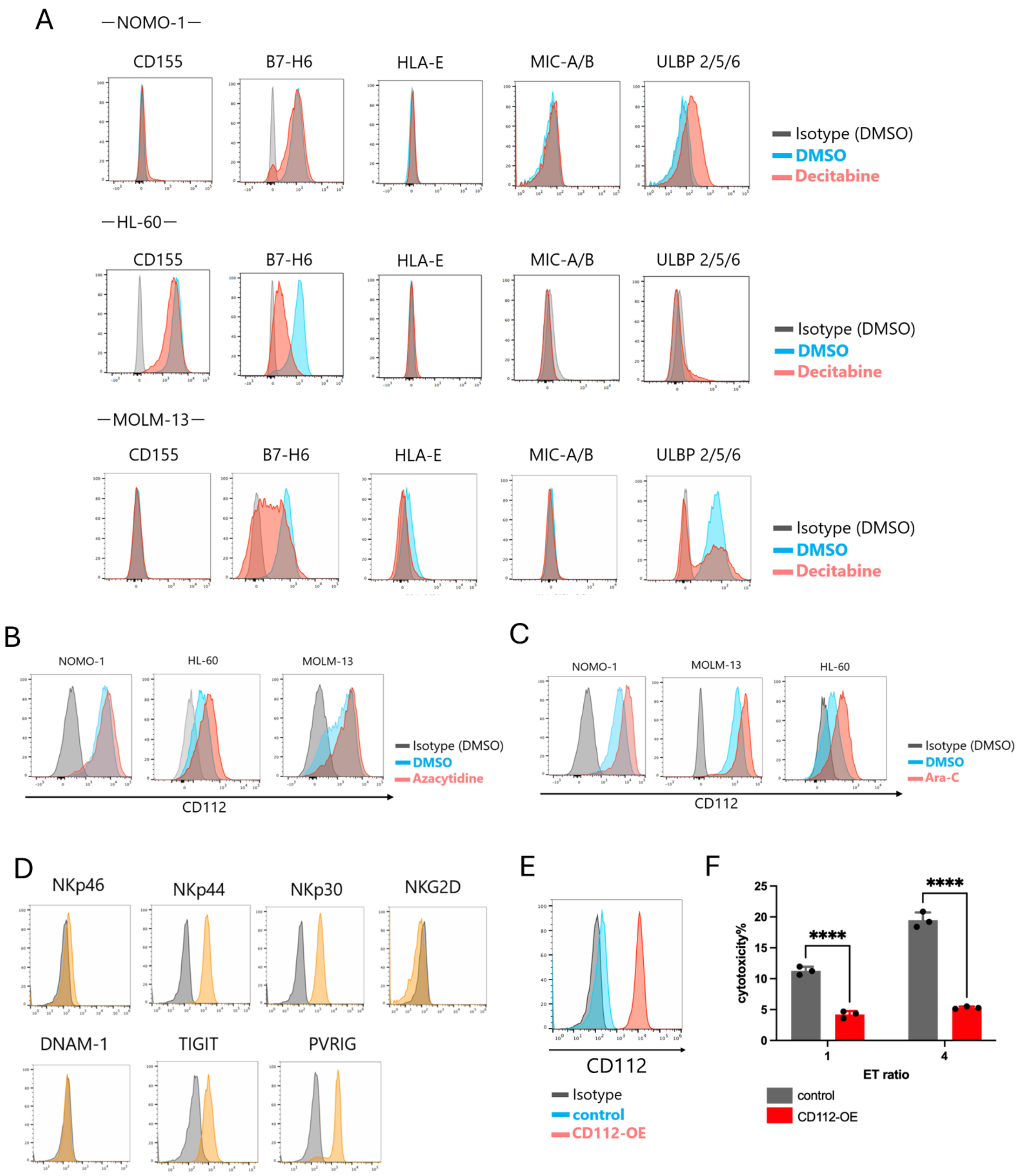

**Supplementary Figure 2. Expression of immunoregulatory molecules in drug-treated AML cells and NK-92 cells**

(A) Representative flow cytometry histograms showing the surface expression of the indicated immunoregulatory molecules on NOMO-1, HL-60, and MOLM-13 cells following treatment with DMSO or decitabine. (B and C) NOMO-1, HL-60, and MOLM-13 cells were treated with DMSO or azacitidine (B) and with DMSO or Ara-C (C) under the following conditions: azacitidine, NOMO-1, 1.0 μM for 48 h; HL-60, 1.0 μM for 72 h; and MOLM-13, 2.5 μM for 48 h; Ara-C, NOMO-1, 1.0 μM for 48 h; HL-60, 3.0 μM for 72 h; and MOLM-13, 0.01 μM for 48 h. CD112 surface expression was subsequently analyzed by flow cytometry. (D) The surface expression of the indicated immunoregulatory receptors on NK-92 cells was analyzed by flow cytometry. Gray histograms indicate the isotype controls, and orange histograms indicate the expression of the respective receptors. (E) CD112 surface expression on HL-60 cells transduced with an empty pMYs-IP vector (Control) or a CD112-expression vector (CD112-OE) was analyzed by flow cytometry following puromycin selection. (F) HL-60 cells transduced with an empty pMYs-IP vector (Control) or a CD112-expression vector (CD112-OE) were co-cultured with NK-92 cells at effector-to-target (E:T) ratios of 1:1 (left) and 4:1 (right) for 4 h at 37°C. Data shown are representative of three independent experiments. Data are presented as the mean ± SD. Data were analyzed by two-way ANOVA followed by a multiple-comparisons test. ns, not significant; ****$P < 0.0001$.

#### Supplementary Figure 3

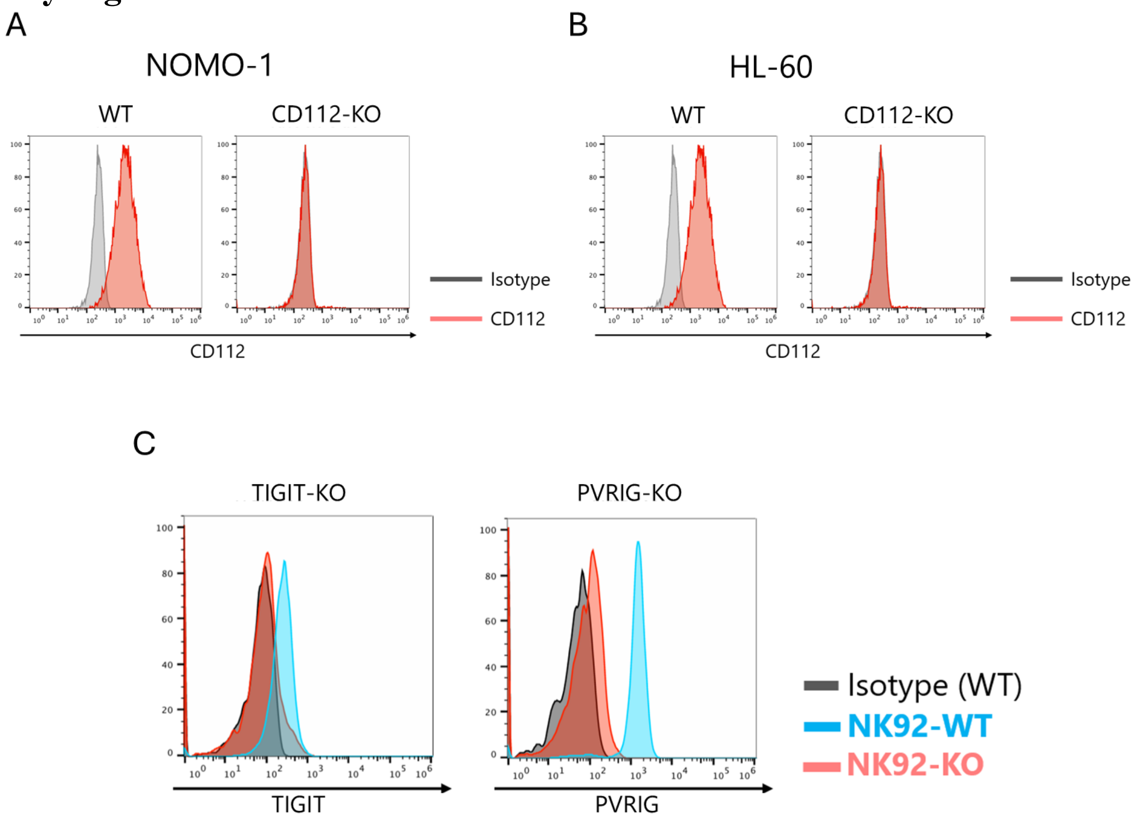


**Supplementary Figure 3. Validation of CRISPR–Cas9-mediated knockout cells**

(A and B) CD112 surface expression on wild-type (WT) and CD112-knockout NOMO-1 (A) and HL-60 (B) cells was analyzed by flow cytometry following puromycin selection. (C) TIGIT surface expression on WT and TIGIT-knockout NK-92 cells was analyzed by flow cytometry following puromycin selection. PVRIG-knockout NK-92 cells were further enriched by sorting for cells with absent or very low PVRIG surface expression, and PVRIG surface expression on WT and sorted PVRIG-knockout NK-92 cells was analyzed by flow cytometry. Data shown are representative of three independent experiments.

### Supplementary Figure 4

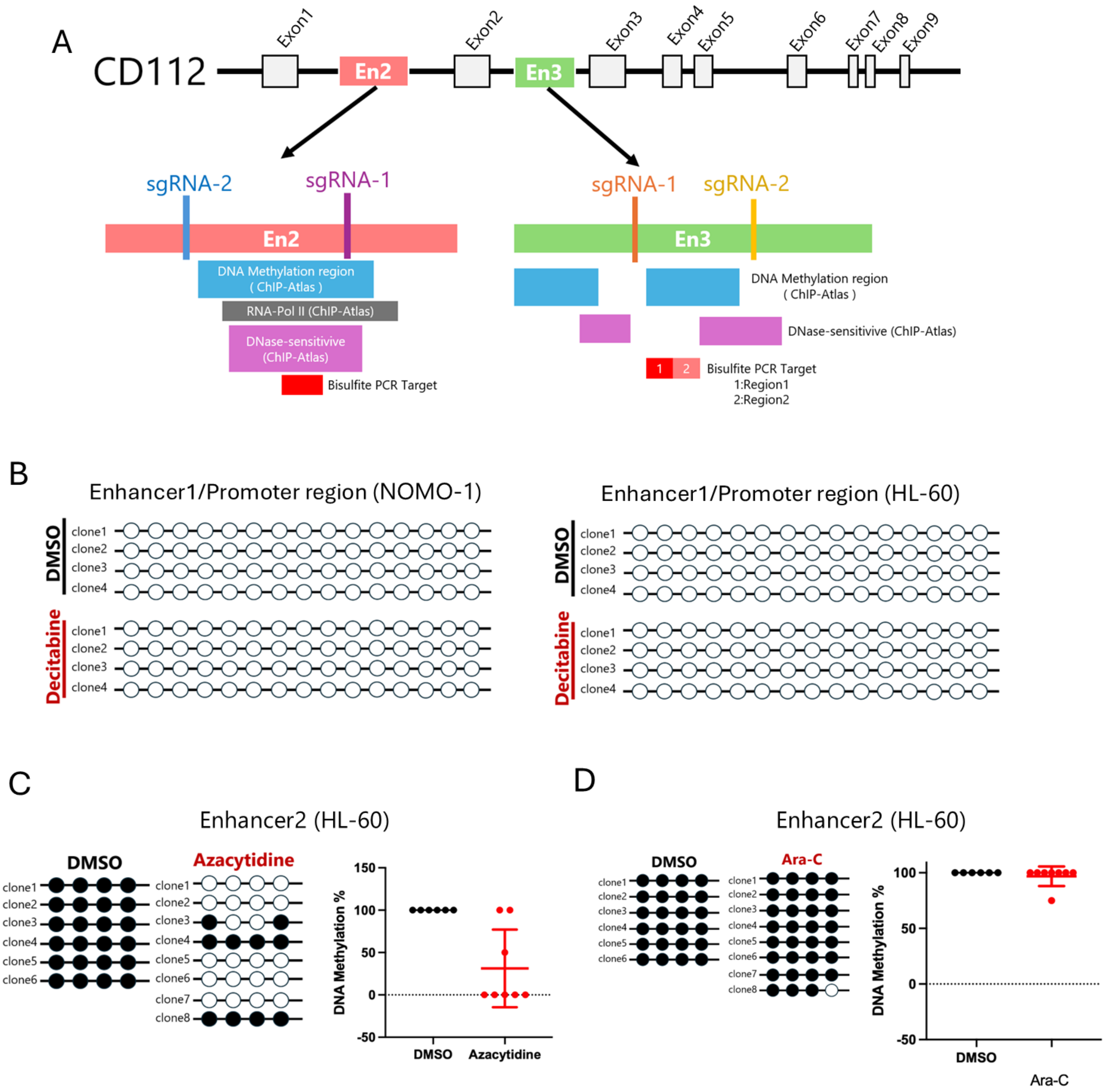

**Supplementary Figure 4. DNA methylation analysis of CD112 regulatory regions**

(A) Schematic illustration of the CD112 gene showing the positions of the bisulfite PCR target regions within enhancers 2 and 3 (En2 and En3), candidate DNA methylation sites, the RNA polymerase II-binding region, and DNase I-hypersensitive regions identified using ChIP-Atlas. (B) NOMO-1 and HL-60 cells were treated with decitabine or DMSO, and genomic DNA was subjected to bisulfite sequencing analysis of the CD112 enhancer/promoter region. Lollipop plots show the methylation status of individual CpG sites in each clone. Open and filled circles indicate unmethylated and methylated CpG sites, respectively. (C) HL-60 cells were treated with DMSO or 1 μM azacitidine for 72 h. Genomic DNA from each sample was subjected to bisulfite sequencing analysis of the CD112 enhancer 2 (En2) region. The percentage of methylated CpG sites within the region was calculated from the bisulfite sequencing results and is shown in the right panels. (D) HL-60 cells were treated with DMSO or 3.0 μM Ara-C for 72 h. Genomic DNA from each sample was subjected to bisulfite sequencing analysis of the CD112 enhancer 2 (En2) region.

**Supplementary Figure 5**

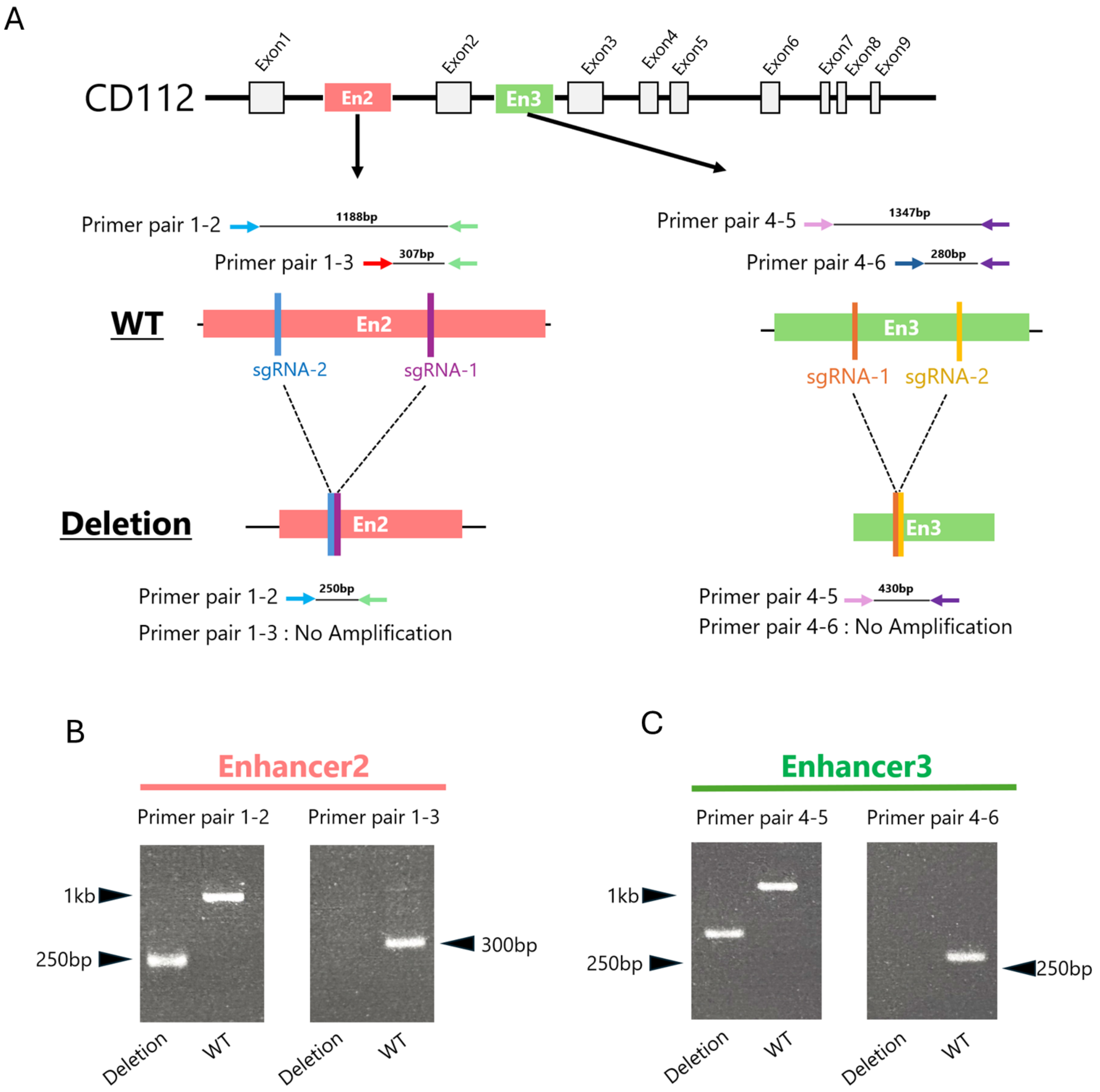

**Supplementary Figure 5. Generation and validation of CD112 enhancer deletion clones**

(A) For each enhancer, two sgRNAs flanking the target region were designed and co-transduced into Cas9-expressing HL-60 cells. Following puromycin and blasticidin selection, single-cell clones were established. Deletion of the target regions was validated by PCR using primer pairs 1–2 and 1–3 for En2 and primer pairs 4–5 and 4–6 for En3. In correctly deleted clones, PCR products amplified with primer pair 1–2 for En2 or primer pair 4–5 for En3 were shorter than those amplified from the corresponding wild-type (WT) alleles, whereas no PCR products were detected using primer pair 1–3 for En2 or primer pair 4–6 for En3. (B and C) En2 (B) and En3 (C) deletion clones were validated by PCR. Deleted clones produced shorter PCR products than WT cells when amplified with primer pair 1–2 for En2 or primer pair 4–5 for En3, whereas no PCR products were detected using primer pair 1–3 for En2 or primer pair 4–6 for En3.

**Supplementary Table 1. Antibodies Used for Flow Cytometric Staining**

| Antibody | Fluorochrome | Clone | Manufacturer | Catalog Number |
|---|---|---|---|---|
| Anti-humanCD112 | PE | TX-31 | Biolegend | 337409 |
| Anti-humanCD155 | PE | SK1L4 | Biolegend | 337609 |
| Anti-humanB7H6 | Alexa Fluor 647 | 1A5 | BD Bioscience | 566733 |
| Anti-humanHLAE | PE | 3D12 | Biolegend | 342603 |
| Anti-humanMICA/B | BV421 | 159227 | BD Bioscience | 749779 |
| Anti-humanULBP2/5/6 | BV422 | 165903 | BD Bioscience | 748128 |
| Anti-humanDNAM1 | PE | 11A8 | Biolegend | 338312 |
| Anti-humanTIGIT | PE | MBSA43 | invitrogen | 12-9500-42 |
| Anti-humanPVRIG | APC | W16216D | Biolegend | 301506 |
| Anti-humanNKp46 | PE | 9E2 | Biolegend | 221907 |
| Anti-humanNKp44 | PE | P44-8 | Biolegend | 325107 |
| Anti-humanNKp30 | PE | P30-15 | Biolegend | 325207 |
| Anti-humanNKG2D | PE | 1D11 | Biolegend | 320805 |

**Supplementary Table 2. sgRNA Sequences for CRISPR/Cas9-Mediated Gene Knockout**

| Name (sgRNA oligonucleotide) | Sequence (5–3′) |
|---|---|
| human-CD112 sgRNA Forward | CGTCCTCCACCGTGAGCCCG |
| human-CD112 sgRNA Reverse | CGGGCTCACGGTGGAGGACG |
| human-TIGIT sgRNA Forward | CAGGCCTTACCTGAGGCGAG |
| human-TIGIT sgRNA Reverse | CTCGCCTCAGGTAAGGCCTGC |
| human-PVRIG sgRNA Forward | ATGCTCGTGCTCTCGGTGCC |
| human-PVRIG sgRNA Reverse | GGCACCGAGAGCACGAGCAT |
| human-CD112 Enhancer2 deletion sgRNA1 Forward | ACATGCACCACAGTCAGATC |
| human-CD112 Enhancer2 deletion sgRNA1 Reverse | GATCTGACTGTGGTGCATGT |
| human-CD112 Enhancer2 deletion sgRNA2 Forward | AATGGAAGGGATTTGGTAGT |
| human-CD112 Enhancer2 deletion sgRNA2 Reverse | ACTACCAAATCCCTTCCATT |
| human-CD112 Enhancer3 deletion sgRNA1 Forward | TCCAGAGATAACTTCAGGGC |
| human-CD112 Enhancer3 deletion sgRNA1 Reverse | GCCCTGAAGTTATCTCTGGA |
| human-CD112 Enhancer3 deletion sgRNA2 Forward | GAAAGTGCCCTTGACGAAGT |
| human-CD112 Enhancer3 deletion sgRNA2 Reverse | ACTTCGTCAAGGGCACTTTC |

**Supplementary Table 3. Primer sequences used for qPCR**

| Name (RT-qPCR primer) | Sequence (5–3′) |
|---|---|
| human-CD112 RT-qPCR Forward | GAGCAGATGGTGTCACGGTCAC |
| human-CD112 RT-qPCR Reverse | GAGATGGACACTTCAGGAGGGT |
| human-GAPDH RT-qPCR Forward | GAAGGTGAAGGTCGGAGTCA |
| human-GAPDH RT-qPCR Reverse | TTGAGGTCAATGAAGGGGTC |
